# The CTP-binding domain is disengaged from the DNA-binding domain in a co-crystal structure of *Bacillus subtilis* Noc-DNA complex

**DOI:** 10.1101/2022.02.28.481274

**Authors:** Kirill V. Sukhoverkov, Adam S. B. Jalal, David M. Lawson, Tung B. K. Le

**Author notes:** these authors contributed equally to this work (Kirill V. Sukhoverkov, Adam S. B. Jalal).

## Abstract

In *Bacillus subtilis*, a ParB-like nucleoid occlusion protein (Noc) binds specifically to Noc-binding sites (*NBS*) around the chromosome to help coordinate chromosome segregation and cell division. Noc does so by binding to cytidine triphosphate (CTP) to form large membrane-associated nucleoprotein complexes to physically inhibit the assembly of the cell division machinery. The site-specific binding of Noc to *NBS* DNA is a prerequisite for CTP-binding and the subsequent formation of a membrane-active DNA-entrapped protein complex. Here, we solve the structure of a truncated *B. subtilis* Noc bound to *NBS* DNA to reveal the conformation of Noc at this crucial step. Our structure reveals the disengagement between the N-terminal CTP-binding domain and the *NBS*-binding domain of each DNA-bound Noc subunit, this is driven, in part, by the swapping of helices 4 and 5 at the interface of the two domains. Site-specific crosslinking data suggest that this conformation of Noc-*NBS* exists in solution. Overall, our results lend support to the recent proposal that *parS/NBS*-binding catalyzes CTP-binding and DNA-entrapment by preventing the re-engagement of the NTD and DBD from the same ParB/Noc subunit.

## INTRODUCTION

Cells must couple chromosome segregation and division to reproduce efficiently. In Firmicutes, such as *Bacillus subtilis*, the nucleoid occlusion protein Noc contributes to the coordination between chromosome segregation and the initiation of cell division^1–4^. Noc helps direct the assembly of the cell division machinery towards the middle of a dividing cell where the concentration of DNA is the least, thus increasing cell division efficiency^2–4^. Critical to this function of Noc is its ability to recruit chromosomal DNA to the cell membrane to form large Noc-DNA-membrane complexes which inhibit the FtsZ formation over the nucleoid and/or to corral the FtsZ ring towards the mid-cell position^5,6^. Noc is a paralog of a chromosome partitioning protein ParB, and is also a CTPase enzyme that binds cytidine triphosphate (CTP) to form a protein clamp that can slide and entrap DNA^6–8^. Apo-Noc first binds to nucleate on 16-bp *NBS* (Noc-binding site) sites scattering along the chromosome^6,9,10^. The nucleation at *NBS* promotes CTP-binding and the subsequent engagement of N-terminal domains from opposing subunits of a Noc homodimer to form a clamp-closed complex that can escape from *NBS* to slide and spread to the neighboring DNA while still entrapping DNA^6^. The DNA-entrapped Noc-CTP complexes are also active at binding to the cell membrane due to the liberation of a 10-amino-acid membrane-targeting amphipathic helix^6^. As a result, Noc-CTP brings the entrapped chromosomal DNA close to the cell membrane to form large Noc-DNA-membrane complexes that are inhibitory to the assembly of nearby cell division machinery^5,6^.

Previously we solved two X-ray crystallography structures of the CTP-binding domain and DNA-binding domain of a *Geobacillus thermoleovorans* Noc to better understand the molecular mechanism of this protein family^6^. Nevertheless, it remains unclear how the Noc-*NBS* binding event mechanistically promotes the N-terminal domain engagement to form a closed-clamp Noc. To investigate further, in this study, we solve a structure of a *B. subtilis Noc-NBS* DNA complex to reveal the conformation of a nucleating Noc. Through comparisons to other available structures of Noc, and its paralog ParB, and by in-solution site-specific crosslinking, we provide evidence for the extended conformation of nucleating Noc.

## RESULTS

### Co-crystal structure of *B. subtilis* Noc with *NBS* DNA reveals that the N-terminal CTP-binding domain of each Noc subunit is disengaged from its DNA-binding domain

To gain insight into the nucleating state of Noc, we sought to determine a co-crystal structure of a Noc-*NBS* complex from *B. subtilis*. After screening several constructs with various lengths of Noc and *NBS*, we solved a 2.9 Å resolution crystal structure of *B. subtilis* NocΔCTD in complex with 16-bp *NBS* DNA duplex (see Materials and Methods) (Table 1). The NocΔCTD variant lacks the 41-amino-acid C-terminal domain (CTD) responsible for Noc dimerization (Figure 1A-B)^5,6^. The asymmetric unit contains two copies of NocΔCTD bound to a single 16-bp *NBS* DNA duplex (Figure 1B).

**Figure 1.**
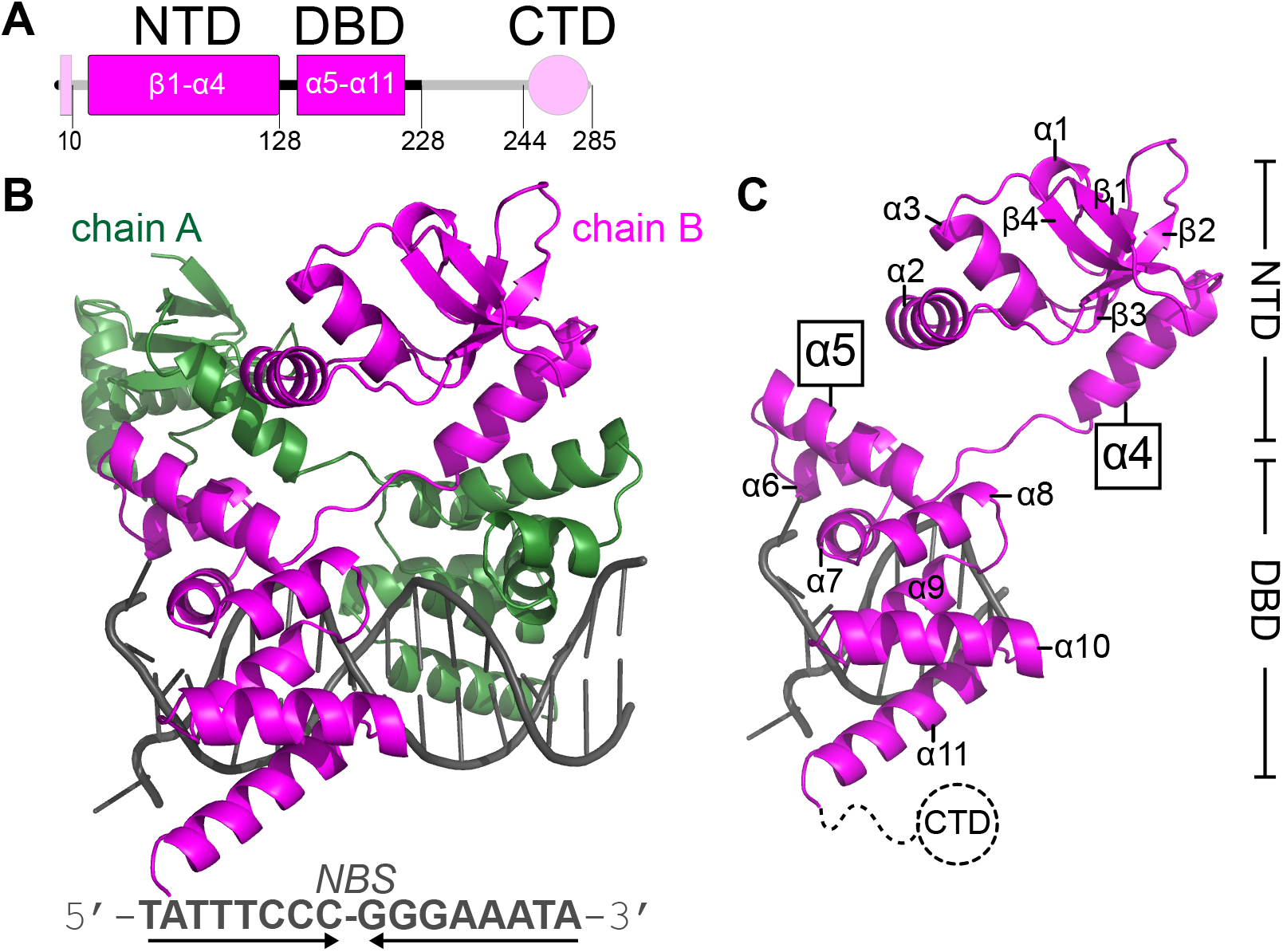
Co-crystal structure of *B. subtilis* Noc with *NBS* DNA reveals that the N-terminal CTP-binding domain of each Noc subunit is disengaged from its DNA-binding domain. **(A)** The domain architecture of *B. subtilis* Noc: the 10-amino-acid N-terminal membrane-targeting peptide, the N-terminal CTP-binding domain (NTD), the central DNA-binding domain (DBD), and the C-terminal domain (CTD). Segments of Noc that are not observed in the structure of NocΔCTD-*NBS* DNA are shown in pale magenta. **(B)** Co-crystal structure of two NocΔCTD subunits (chain A: dark green, chain B: magenta) bound to a 16-bp *NBS* DNA duplex (grey). The nucleotide sequence of a 16-bp *NBS* is shown below the crystal structure, converging arrows indicate that *NBS* is palindromic. **(C)** The structure of chain B of NocΔCTD (magenta) bound to an *NBS* half-site with key features such as the swinging-out helices α4-α5 highlighted.

**Table 1.** X-ray data collection, processing, and refinement statistics.

|  |  |
| --- | --- |
| *Data collection* |  |
| Diamond Light Source beamline | I04 |
| Wavelength (Å) | 0.979 |
| Detector | Eiger2 XE 16M |
| Resolution range (Å) | 70.27 – 2.90 (3.08 – 2.90) |
| Space Group | *P2<sub>1</sub>2<sub>1</sub>2<sub>1</sub>* |
| Cell parameters (Å) | *a* = 70.5, *b* = 99.3, *c* = 99.4 |
| Total no. of measured intensities | 164414 (27117) |
| Unique reflections | 16090 (2546) |
| Multiplicity | 10.2 (10.7) |
| Mean *I*/$\sigma$(*I*) | 11.7 (1.2) |
| Completeness (%) | 100.0 (100.0) |
| *R*<sub>merge</sub><sup>a</sup> | 0.097 (1.976) |
| *R*<sub>meas</sub><sup>b</sup> | 0.102 (2.076) |
| *CC*<sub>1/2</sub><sup>c</sup> | 0.999 (0.691) |
| Wilson *B* value (Å<sup>2</sup>) | 85.6 |
| *Refinement* |  |
| Resolution range (Å) | 70.27 – 2.90 (2.98 – 2.90) |
| Reflections: working/free<sup>d</sup> | 15216/1101 |
| *R*<sub>work</sub><sup>e</sup> | 0.230 (0.407) |
| *R*<sub>free</sub><sup>e</sup> | 0.278 (0.419) |
| Ramachandran plot:<br>favored/allowed/disallowed<sup>f</sup> (%) | 96.5/3.5/0 |
| R.m.s. bond distance deviation (Å) | 0.003 |
| R.m.s. bond angle deviation (°) | 1.18 |
| Mean *B* factors: protein/DNA/overall (Å<sup>2</sup>) | 123/86/117 |
| **PDB accession code** | **7OL9** |

Values in parentheses are for the outer resolution shell.
<sup>a</sup> $R_{\text{merge}} = \sum_{hkl} \sum_i |I_i(hkl) - \langle I(hkl) \rangle| / \sum_{hkl} \sum_i I_i(hkl)$ .
<sup>b</sup> $R_{\text{meas}} = \sum_{hkl} [N/(N-1)]^{1/2} \times \sum_i |I_i(hkl) - \langle I(hkl) \rangle| / \sum_{hkl} \sum_i I_i(hkl)$ , where $I_i(hkl)$ is the *i*th observation of reflection *hkl*, $\langle I(hkl) \rangle$ is the weighted average intensity for all observations *i* of reflection *hkl* and *N* is the number of observations of reflection *hkl*.
<sup>c</sup> *CC*<sub>1/2</sub> is the correlation coefficient between symmetry equivalent intensities from random halves of the dataset.
<sup>d</sup> The dataset was split into "working" and "free" sets consisting of 95 and 5% of the data respectively. The free set was not used for refinement.
<sup>e</sup> The R-factors *R*<sub>work</sub> and *R*<sub>free</sub> are calculated as follows: $R = \sum (|F_{\text{obs}} - F_{\text{calc}}|) / \sum |F_{\text{obs}}|$ , where *F*<sub>obs</sub> and *F*<sub>calc</sub> are the observed and calculated structure factor amplitudes, respectively.
<sup>f</sup> As calculated using MolProbity<sup>29</sup>

Each NocΔCTD subunit consists of an N-terminal CTP-binding domain (NTD) (helices α1 to 4 and sheets β1 to 4) and a DNA-binding domain (DBD) (helices α5 to 11) (Figure 1C). The electron density for the first 27 amino acids that contains the membrane-targeting peptide was poorly resolved, and thus this region was absent from the model (Figure 1A). Each NocΔCTD subunit is bound to a half *NBS* site, the *NBS* DNA adopts a conformation whereby in one strand the 5’ base was flipped out, and in the other, the 3’ base was flipped out, enabling a sticky-ended interaction (with a one-base overhang) between the duplexes in adjacent asymmetric units (Figure 1B and Supplementary Figure 1). We previously solved a structure of only the DBD of Noc with *NBS* (2.23Å, PDB: 6Y93) to elucidate the molecular basis for *NBS*-binding specificity^10^. Given that the conformation of the DBD and the core *NBS* site are similar between the previous structure and the structure in this work (root-mean-square deviation RSMD = 0.46 Å), we describe the conformation of the NTD in-depth here instead. By structural alignment of the two NocΔCTD subunits, we noted that the DBD and helices α4-5 are highly similar (RSMD = 0.27 Å) while the rest of the NTD (β1-β4) is orientated in a different direction (approx. 30° apart, owing to the flexible loop in between α4 and β4) (Figure 2A). The multiple alternative orientations at the NTD is likely a common feature of all nucleating ParB family proteins, including Noc. This was the case for the NTD of *Caulobacter crescentus* ParB bound to *parS* DNA^11^, and is also evidential from the superimposition of the *B. subtilis* NocΔCTD-*NBS* structure onto that of ParBΔCTD-*parS* from *Helicobacter pylori* and *C. crescentus* (Figure 2B)^11,12^. Multiple alternative conformations of nucleating ParB/Noc family members suggest flexibility at the N-terminal CTP-binding domain.

**Figure 2.**
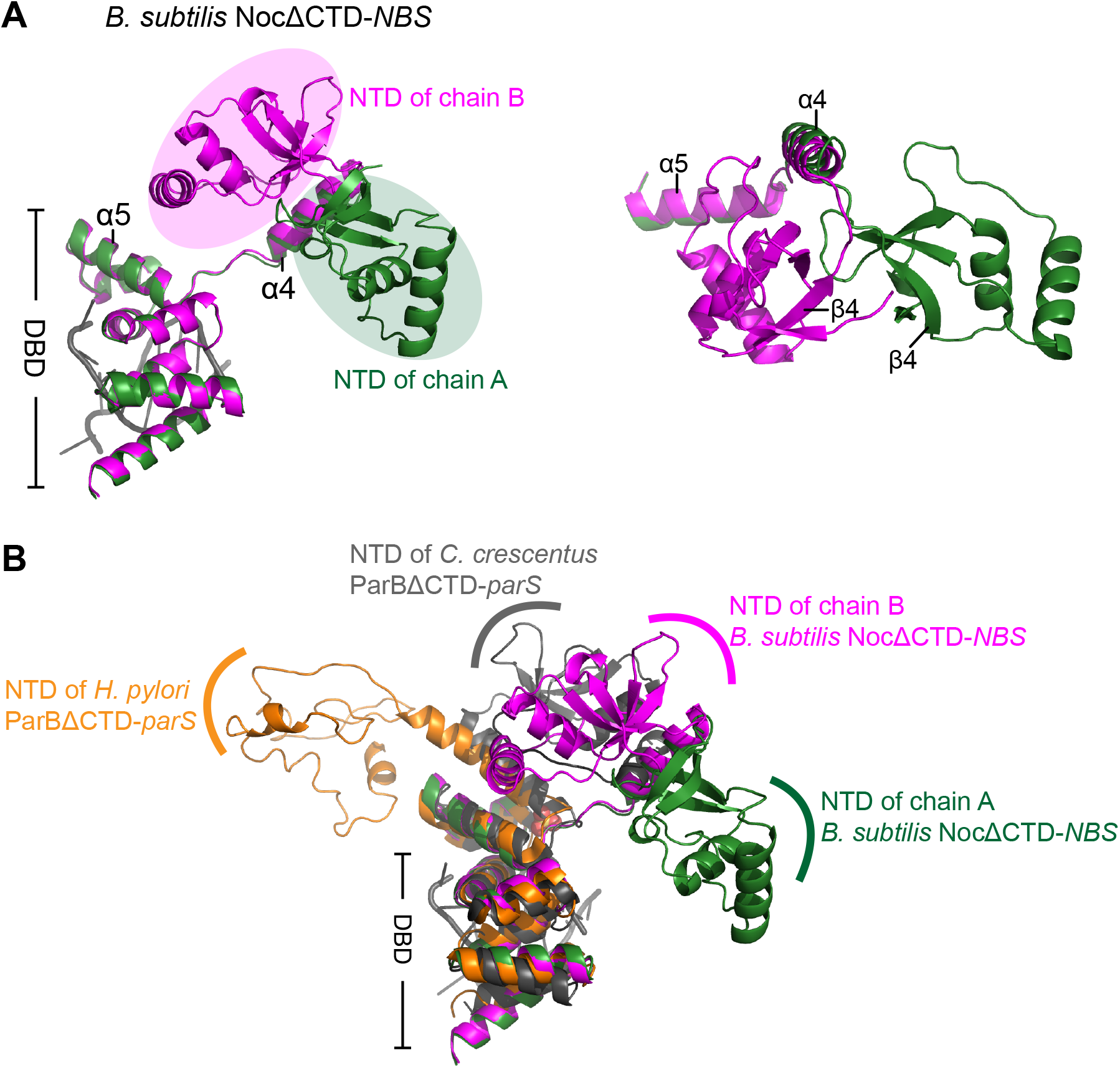
Co-crystal structure of *B. subtilis* NocΔCTD-*NBS* shows alternative orientations at the N-terminal domain (NTD) of Noc. **(A)** (left panel) Superimposition of chain A and chain B of NocΔCTD shows the different orientations of the NTD. (right panel) the top-down view of the superimposition of NocΔCTD subunits shows the majority of the NTD orientates ~30° apart; part of the DBD (from α6 to α11) was omitted for clarity. Half of the 16-bp palindromic *NBS* site (grey) is shown. **(C)** Structural superimposition of *B. subtilis* NocΔCTD-*NBS* upon other available DNA-bound ParB structures highlights the variation in the orientation of the NTD.

The most notable feature of the NocΔCTD-*NBS* structure is the disengagement of the NTD and DBD (Figure 1C), which is likely driven by the swinging-out conformation of α4-α5 (Figure 3A-B). Helices α4 and α5 from the same NocΔCTD subunit are not packed together, instead α4 swings outwards by approx. 100° to pack against α5’ from the adjacent NocΔCTD subunit (Figure 3A). This swinging-out conformation has not been observed in the previous structures of DNA-bound *C. crescentus* or *H. pylori* ParBΔCTD, of *Thermus thermophilus* ParBΔCTD-apo, or *G. thermoleovorans* NocΔCTD-apo (Figure 3B and Supplementary Figure 2)^6,11–13^. In previous structures of apo- or DNA-bound ParB/Noc, the equivalent helix α4 consistently folds back to pack with α5 from the same protein subunit (the folding-back conformation) (Figure 3B and Supplementary Figure 2). The swinging-out conformation of helices α4-5 is often associated with the nucleotide-bound state of ParB/Noc instead (Figure 3B and Supplementary Figure 2)^6,7,11,14^. It has been suggested that CTP-binding most likely facilitates the swinging-out conformation of ParB/Noc since nucleotides have been observed to make numerous contacts to both the equivalent α4 and the α4-α5 connecting loop in various ParB proteins^7,8,11,14^. The observation of a swinging-out conformation in DNA-bound Noc is therefore surprising, given that CTP was not included in the crystallization drop and that CTP-binding is incompatible with high-affinity binding at the nucleation site *NBS*^6^. We reason that the swinging-out conformation might be thermodynamically possible in the DNA-bound nucleating ParB/Noc, and that CTP-binding, instead of facilitating, further stabilizes the swinging-out conformation.

**Figure 3.**
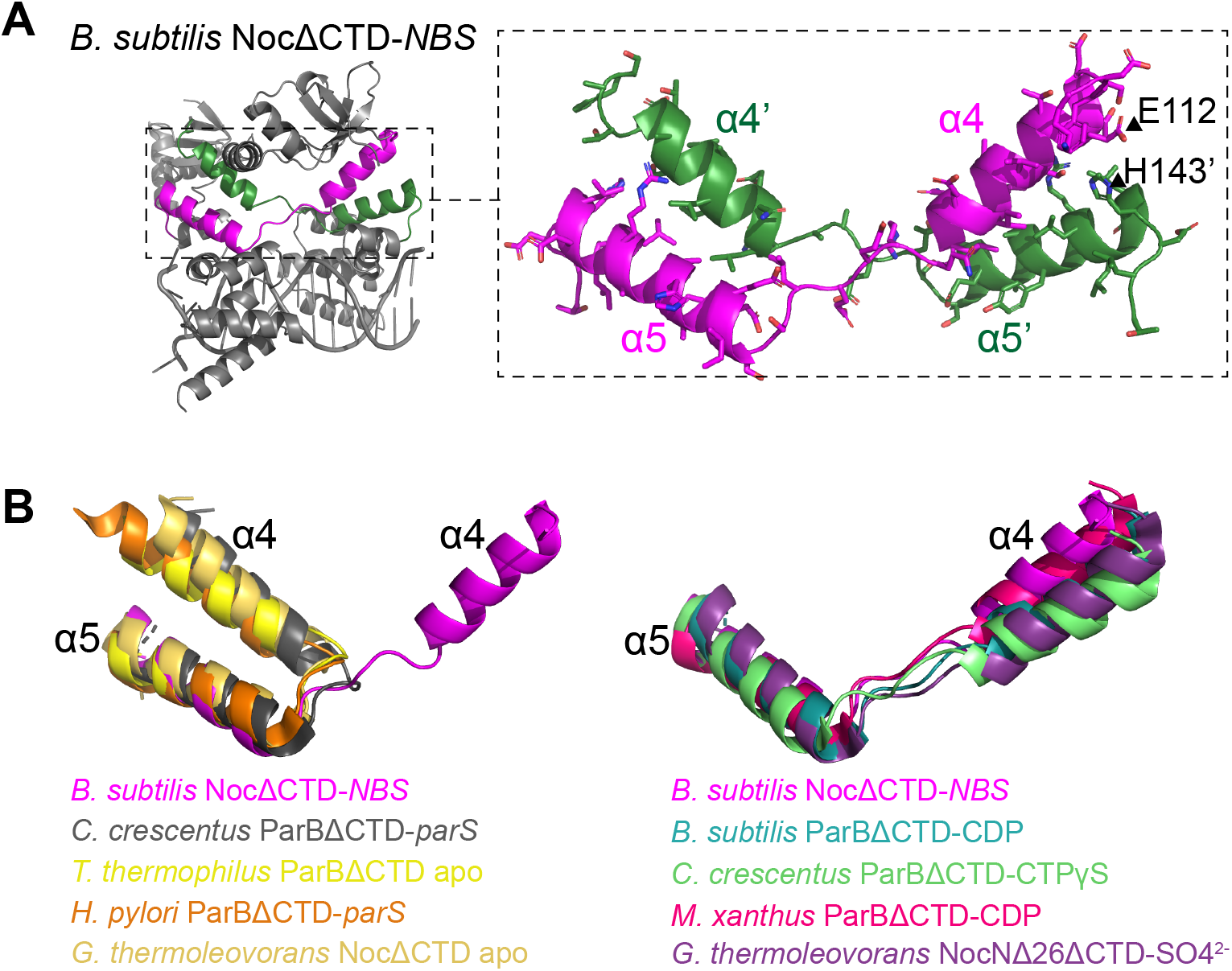
Helices α4-α5 from the *B. subtilis* NocΔCTD-*NBS* complex adopt a swinging-out conformation. **(A)** Co-crystal structure of *B. subtilis* NocΔCTD with *NBS* (left panel) with the pairs of swapping helices (α4-α5, and α4’-α5’ for the opposite subunit) highlighted in magenta and dark green, respectively (right panel). Amino acid side chains of α4-α5 and α4’-α5’ are shown in stick representation to illustrate the packing between helices from opposite Noc subunit. The positions of residues E112 (on α4) and H143 (on α5), which were substituted for cysteine in a crosslinking assay (Figure 4), are also shown. **(B)** Superimposition of the helices α4-α5 from *B. subtilis* NocΔCTD-*NBS* complex upon the equivalent pair of helices in other apo-ParB/Noc or DNA-bound ParB structures (left panel), or upon the equivalent pair of helices in other nucleotide-bound ParB/Noc structure (right panel). The *G. thermoleovorans* NocNΔ26ΔCTD-SO4^2-^ structure is thought to represent the conformation of Noc in a nucleotide-bound state as the sulfate anion from crystallization solution occupies a position equivalent to that of the β-phosphate moiety of CTP^6^.

### Site-specific cysteine-cysteine crosslinking suggests the swinging-out conformation of Noc-*NBS* in solution

To test if the swinging-out conformation of α4-α5 is possible in *NBS*-bound Noc in solution, we employed site-specific chemical crosslinking with the cysteine-specific bismaleimide compound BMOE^15^. Based on the structures of apo-NocΔCTD^6^ and NocΔCTD-*NBS*, we engineered a dual cysteine substitution at E112 and H143 on an otherwise cysteine-free *B. subtilis* Noc to create a Noc (E112C H143C) variant (Figure 3A). In the folding-back conformation where helices α4 and α5 from the same Noc subunit pack together, crosslinking of E112C to H143C would generate an intramolecular crosslinked species (Noc IntraXL), while a swinging-out conformation would give rise to intermolecularly crosslinked species (a singly-crosslinked Noc InterXL and a doubly-crosslinked Noc Inter2XL) which are twice the theoretical molecular weight of a Noc monomer (Figure 4A). Crosslinking of apo-Noc (E112C H143C) only resulted in a prominent band that migrated faster in a denaturing acrylamide gel than noncrosslinked protein (Figure 4B, lane 1 vs. 2), this is most likely a Noc IntraXL species. Little of Noc InterXL or Inter2XL species was observed (~4.4% crosslinked fraction) suggesting that the swinging-out conformation is unfavored in apo-Noc (Figure 4B, lane 1 vs. lane 2). The addition of only CTP did not promote the swinging-out conformation noticeably (Figure 4B, lane 2, ~4.4% vs. lane 4, ~8.7% crosslinked fraction). The singly (InterXL) and the doubly (Inter2XL) crosslinked species appeared more prominently when *NBS* only (Figure 4B, lane 2, ~4.4% vs. lane 3, ~19.3% crosslinked fraction) or *NBS* + CTP were preincubated with Noc (Figure 4B, lane 2, ~4.4% vs. lane 5, ~31.5% crosslinked fraction). The InterXL/2XL fraction further increased when *NBS* was used in a molar excess to Noc (E112C H143C) (Supplementary Figure 3). We were able to assign different bands to either being InterXL or Inter2XL by performing crosslinking reactions of Noc (E112C H143C) + *NBS* + CTP with an increasing concentration of the BMOE crosslinker (Figure 4C). The assumption is that a singly-crosslinked InterXL preferably forms at a lower concentration of a crosslinker. Overall, our result here suggests that the swinging-out conformation of α4-5 is possible in solution and is promoted when Noc is bound to the *NBS* DNA.

**Figure 4.**
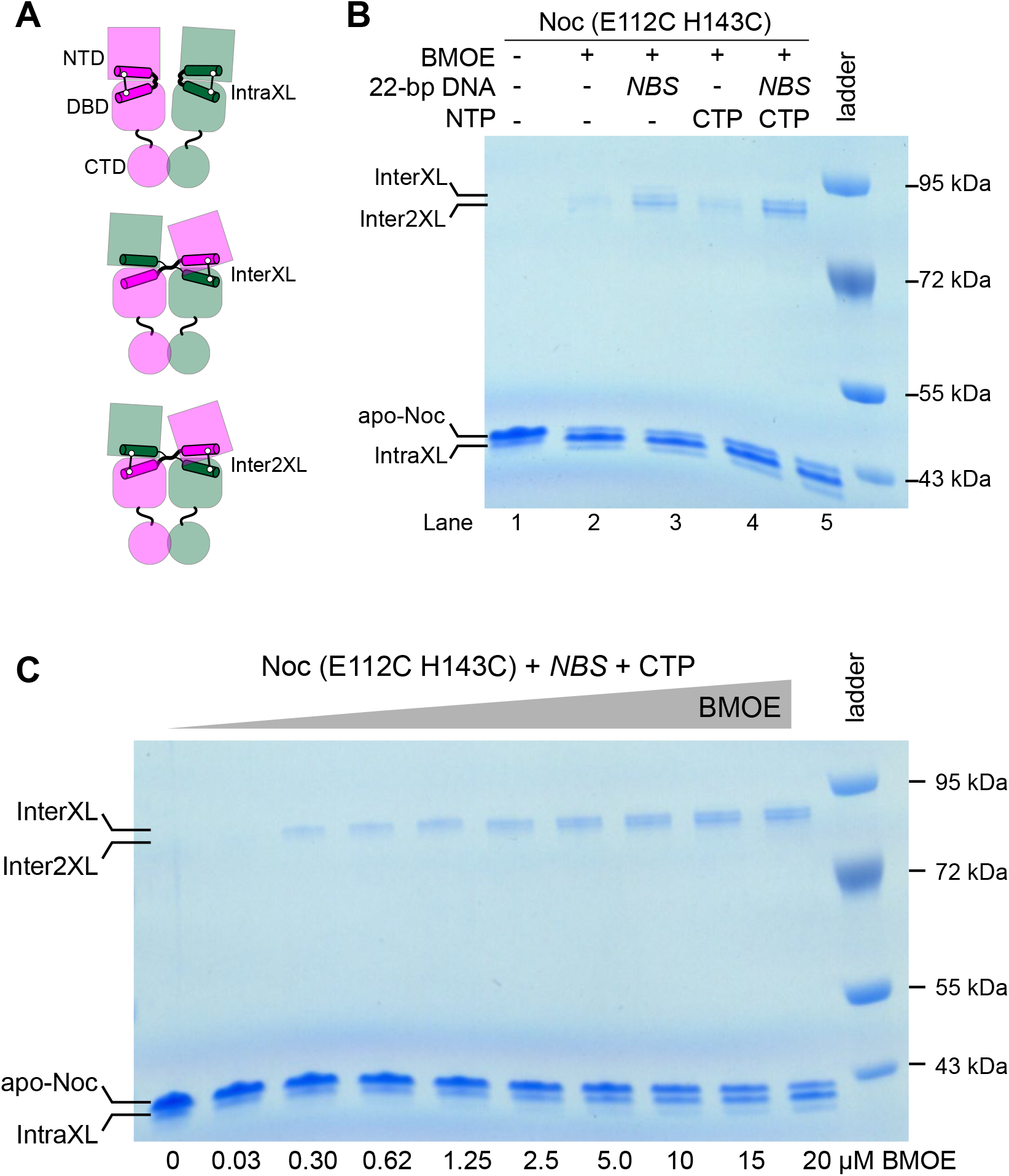
Site-specific crosslinking of *B. subtilis* Noc (E112C H143C) suggests that the swinging-out conformation of Noc-*NBS* exists in solution. **(A)** Schematic diagram of crosslinked species: IntraXL and InterXL/Inter2XL denote crosslinked species formed between α4 and α5 from either the same Noc subunit or from opposing subunits, respectively. Inter2XL represents a double crosslinking between α4 and α5’, and between α4’ and α5. **(B)** Crosslinking of Noc (E112C H143C) in the presence or absence of *NBS* and CTP and combinations thereof. Crosslinked species were resolved on an acrylamide gel and stained with Coomassie. Each crosslinking experiment was run in triplicate. **(C)** Crosslinking of *B. subtilis* Noc (E112C H143C) + CTP + *NBS* with an increasing concentration of BMOE. Crosslinked species were resolved on an acrylamide gel and stained with Coomassie. The top-most band appeared first as the concentration of BMOE is increasing from left to right, and thus it is thought to represent the singly crosslinked InterXL species. The doubly crosslinked Inter2XL presumably is more compacted and migrated faster than the InterXL on a denaturing acrylamide gel. Each crosslinking experiment was run in triplicate.

## DISCUSSION

In *B. subtilis, noc* resulted from *parB* via a gene duplication and neo-functionalization event^10,16^, and both Noc and ParB are CTP-dependent molecular switches^7,8,17–20^. CTP-binding switches nucleating ParB/Noc (bound at a high-affinity *parS/NBS* site) from an open-clamp conformation (Figure 5A-B) to a closed-clamp conformation that can escape from *parS/NBS* to slide to neighboring DNA while still entrapping DNA (Figure 5C)^6,7,14,17,18^. The closed-clamp conformation is possible due to the new dimerization interface between the two adjacent N-terminal CTP-binding domains of ParB/Noc (the so-called NTD-NTD engagement, Figure 5C)^6,7,14,17^. Here, our NocΔCTD-*NBS* structure represents an open-clamp conformation because there is no protein-protein contact between the majority of two adjacent NTDs of Noc, except for the swapping helices α4 and α4’ (Figure 1B and Figure 5B).

**Figure 5.**
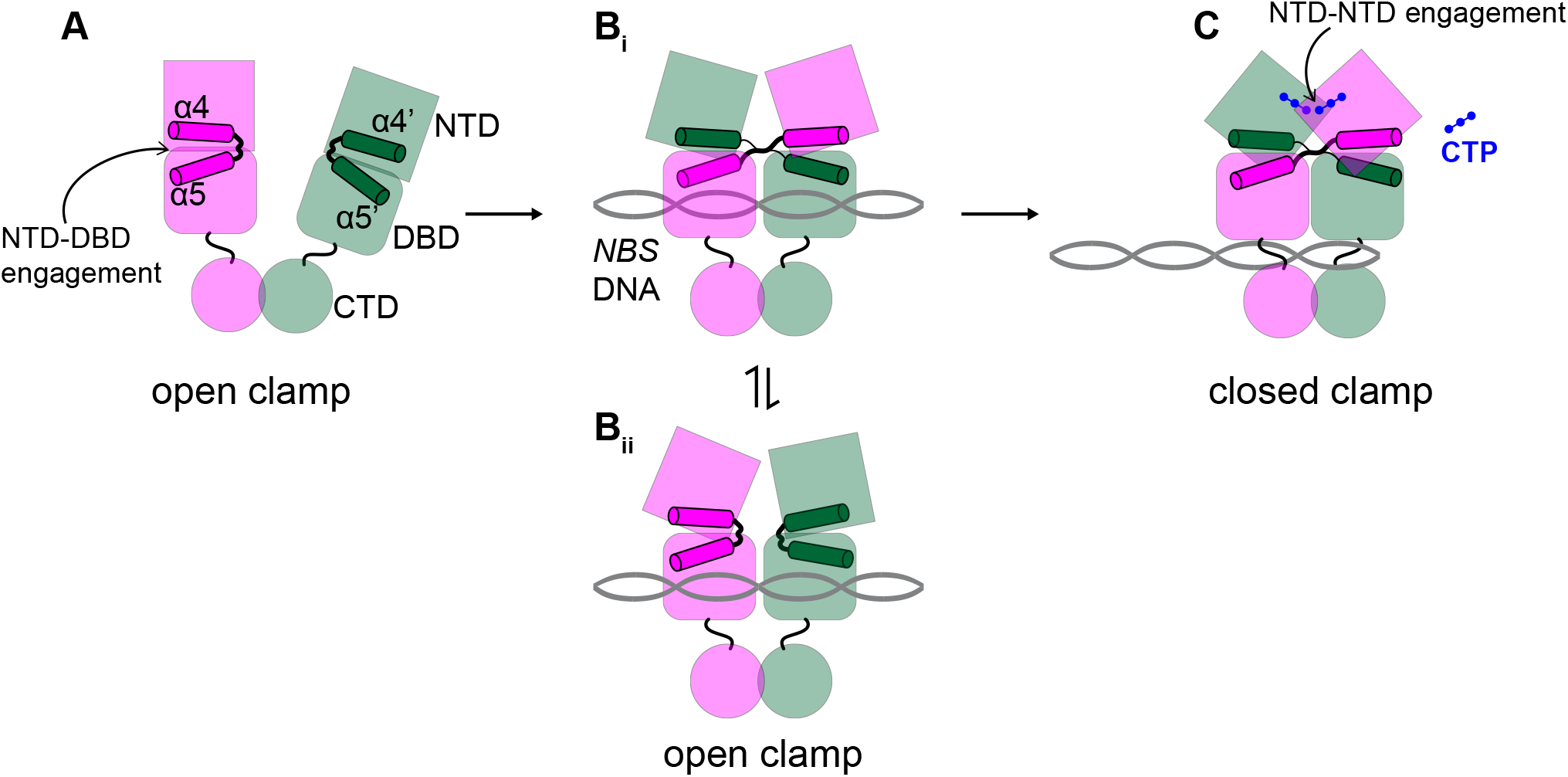
A possible model of different conformations of *B. subtilis* Noc. In apo-Noc (A), helices α4 and α5 from the same Noc subunit (magenta or dark green) likely pack together i.e. the folding-back conformation^6^. In *NBS*-bound Noc, helices α4 and α5 from the same Noc subunit are not packed together, instead α4 swings outwards to pack against α5’ from the adjacent Noc subunit i.e. the swinging-out conformation (B_i_). As the result, the NTD and the DBD from the same Noc subunit disengage from each other, and there is no protein-protein contact between the majority of two adjacent NTDs of Noc. The proximity of adjacent Noc subunits and the restriction in movement by a DNA-bound DBD may increase the likelihood of helix swapping, this might contribute to promoting the NTD-NTD engagement upon CTP-binding (C). It is possible that helices α4 and α5 of *NBS*-bound Noc also take up a folding-back conformation (B_ii_). In this case, NTDs of adjacent subunits of Noc likely adopt different orientations to avoid possible clashing between two adjacent protein subunits^15^, the difference in orientations of opposing NTDs has been observed in the co-crystal structures of ParBΔCTD-*parS* from *C. crescentus* and *H. pylori*^11,12^.

It has been observed that, without *parS/NBS*, CTP is unable to efficiently promote the NTD-NTD engagement to close the ParB/Noc clamp^6,7,14,17^. To rationalize this phenomenon, Antar *et al* (2021) noted that two ParB subunits would not be able to occupy a *parS* site if they were to adopt a conformation similar to apo-ParB (in which the NTD and the DBD of the same ParB subunit fold back on each other) because of a severe clash between opposing ParB subunits^15^. Antar *et al* (2021) proposed that, to avoid this potential clash, the NTD and the DBD from each *parS*-bound ParB must be untethered/disengaged from each other^15^. The DBD-NTD disengagement later favors the two opposing NTDs to dimerize in the presence of CTP to form a clamp-closed complex^15^. In sum, *parS* serves a catalyst in a reaction that favors the formation of the product (the closed clamp) by inhibiting the reversion to the substrate (the open clamp apo-ParB). Our structure of DNA-bound Noc here lends support to this hypothesis because the conformation of the DNA-bound Noc subunit is drastically different from that of apo-Noc, especially with the swinging-out helices α4-α5 disengaging the NTD and DBD from each other (Figure 5Bi). It is possible that *NBS*-bound Noc might exist as an ensemble of states with helices α4-α5 in either a folding-back (Figure 5Bii) or a swinging-out conformation (Figure 5Bi), and that the NocΔCTD-*NBS* structure here represents a snapshot of this dynamic process. The swinging-out conformation of α4-α5 might be rare in solution, given that the crosslinking reaction of Noc (E112C H143C) + *NBS* produced IntraXL as the major species. Nevertheless, the proportion of InterXL and Inter2XL increased substantially when *NBS* (Figure 4B, lane 3) is included in comparison to apo-Noc only (lane 2) or Noc + CTP only conditions (lane 4). Moreover, the proportion of InterXL and Inter2XL also increased when *NBS* was added in excess (Supplementary Figure 3). The proximity of adjacent Noc subunits and the restriction in movement by a DNA-fixated DBD may increase the likelihood of swapping helices α4-α5 in Noc. This might in part contribute to further promoting the NTD-NTD engagement upon CTP-binding (Figure 5C), and might additionally explain how *NBS* serves as a catalysis for NTD-NTD engagement and thus clamp closure for a ParB-like protein Noc. However, it is also worth noting that ParB, in the presence of *parS*, does not undergo α4-α5 swapping as readily as Noc-*NBS* (S. Gruber, personal communication)^15^. It is still unclear why this is the case and how it is related to the biological functions of ParB vs. Noc, but it explains why helices α4-α5 in all previous X-ray crystallography structures of ParB-*parS* complex are all in the folding-back conformation^11,12^.

## ACKNOWLEDGEMENTS

This study was funded by the Royal Society University Research Fellowship Renewal (URF\R\201020 to T.B.K.L) and BBSRC (BBS/E/J/000PR9791 to the John Innes Centre). K.V.S is supported by Wellcome grant (221776/Z/20/Z). A.S.B.J’s PhD studentship was funded by the Royal Society (RG150448). We thank Diamond Light Source for access to beamline I04 under proposal MX18565. We thank Clare Stevenson for assistance with X-ray crystallography. We thank S. Gruber, H. Antar, and T. McLean for discussions and comments on the manuscript.

## MATERIALS AND METHODS

### Plasmid and strain construction

#### Construction of pET21b:: Bacillus subtilis NocΔCTD-his_6_ and pET21b::noc (E112C H143C)-his_6_

The coding sequence of a 41-amino-acid C-terminally truncated *B. subtilis* Noc was amplified by PCR using a forward primer (aactttaagaaggagatatacatatgaagcattcattctctcgtttcttc) and a reverse primer (gtggtgctcgagtgcggccgcaagcttatctctgctgaatgctttgcgtctc), and pET21b::*B. subtilis* Noc-his_6_^6^ as template. The resulting PCR product was gel-purified and assembled into an NdeI-HindIII-cut pET21b using a 2x Gibson master mix (NEB). Gibson assembly was possible owing to a 23-bp sequence shared between the NdeI-and-HindIII cut pET21b backbone and the PCR amplified fragment. The 23-bp homologous region was introduced during the synthesis of the above primers.

A double-stranded DNA (dsDNA) fragment containing a *B. subtilis noc (E112C H143C)* gene was chemically synthesized (gBlocks, IDT). The gBlocks fragment was assembled into an NdeI-HindIII-cut pET21b using a 2x Gibson master mix to result in *pET21b::noc (E112C H143C)-his_6_*. All plasmids were verified by Sanger sequencing (Eurofins, Germany).

### Protein overexpression and purification

*B. subtilis* NocΔCTD-His_6_ were purified through a 3-column (HisTrap, Heparin, Superdex-75 gel filtration) procedure as described previously^6^. Purified NocΔCTD-His_6_ was stored at −80°C in storage buffer (10 mM Tris-HCl pH 8.0 and 250 mM NaCl) before crystallization.

Noc (E112C H143C)-His_6_ was purified through a 2-column (His-Select Cobalt Affinity Gel, Superdex-200 gel filtration) procedure using the following buffers: buffer A-HisTrap (100 mM Tris-HCl pH 7.4, 300 mM NaCl, 10 mM imidazole, 5% (v/v) glycerol), buffer B-HisTrap (100 mM Tris-HCl pH 7.4, 300 mM NaCl, 500 mM imidazole, 5% (v/v) glycerol), and gel filtration buffer (100 mM Tris-HCl pH 7.4 and 300 mM NaCl). Purified protein was concentrated using an Amicon Ultra-4 10 kDa cut-off spin column, and stored at −80°C in storage buffer (100 mM Tris-HCl pH 7.4, 300 mM NaCl, 10 (v/v) glycerol, and 0.1 mM TCEP).

### *In vitro* crosslinking using a sulfhydryl-to-sulfhydryl crosslinker bismaleimidoethane (BMOE)

Noc (E112C H143C)-His_6_ (4 μM final concentration) was incubated on ice either alone or with 1 mM CTP, or 1 μM 22-bp *NBS* DNA duplex (or with a twofold increasing concentration of *NBS* from 0 to 5 μM), or both in a crosslinking buffer (100 mM Tris-HCl pH 7.4, 130 mM NaCl, 5 mM MgCl_2_) for 10 min. Then, 20 mM DMSO solution of the crosslinking reagent (BMOE, ThermoFisher) was added to the reaction to the final concentration of 2 mM. The mixture was incubated at room temperature for 5 min before the crosslinking reaction was quenched by SDS-PAGE loading dye + β-mercaptoethanol. Samples were heated to 90°C for 10 min before being loaded on 4-12% Bis-Tris polyacrylamide gels (ThermoFisher). Each experiment was triplicated. Polyacrylamide gels were stained in an InstantBlue Coomassie solution (Abcam) and band intensity was quantified using Image Studio-Lite (LICOR Biosciences). Raw gel images were deposited to the Mendeley repository: doi: 10.17632/6sp26rm6zy.1

### Reconstitution of *NBS* DNA for X-ray crystallography

A 16-bp *NBS* DNA fragment (5’-TATTTCCCGGGAAATA-3’) (3.6 mM in buffer containing 10 mM Tris-HCl pH 8.0 and 250 mM NaCl) was heated to 98°C for 5 min before being left to cool at room temperature overnight to form double-stranded *NBS* DNA (final concentration: 1.8 mM).

### Protein crystallization, structure determination, and refinement

*B. subtilis* NocΔCTD-His_6_ (~10 mg/mL) was mixed with the 16-bp *NBS* DNA at a molar ratio of 1:1.2 (protein: DNA) in the gel filtration elution buffer (10 mM Tris-HCl pH 8.0, 250 mM NaCl). Crystallization screens were set up in sitting-drop vapor diffusion format in MRC2 96-well crystallization plates with drops comprised of 0.3 μL precipitant solution and 0.3 μL of protein and incubated at 293 K. After optimization of initial hits, the best crystals of the complex grew in a solution containing 17% (w/v) PEG3350, 0.25 M magnesium acetate and 10% (v/v) sucrose. These were cryoprotected in the crystallization solution supplemented with 20% (v/v) glycerol and mounted in Litholoops (Molecular Dimensions) before flash-cooling by plunging into liquid nitrogen. X-ray data were recorded on beamline I04 at the Diamond Light Source (Oxfordshire, UK) using an Eiger2 XE 16M hybrid photon counting detector (Dectris), with crystals maintained at 100 K by a Cryojet cryocooler (Oxford Instruments). Diffraction data were integrated and scaled using DIALS^21^ via the XIA2 expert system^22^ then merged using AIMLESS^23^ to a resolution of 2.9 Å in space group *P*2_1_2_1_2_1_ with cell parameters of *a* = 70.5, *b* = 99.3, *c* = 99.4 Å. Data collection statistics are summarized in Table 1. Analysis of the likely composition of the asymmetric unit (ASU) suggested that it contained two copies of the 29.5 kDa NocΔCTD monomer plus the 16-bp *NBS* duplex, giving an estimated solvent content of 51%.

The majority of the downstream analysis was performed through the CCP4i2 graphical user interface^24^. For molecular replacement, a template was constructed from the structure of the *B. subtilis* NOC DNA-binding domain (DBD) complexed to an *NBS* duplex (PDB accession code 6Y93)^10^. Initially, PHASER^25^ was run using the protein and DNA components of this entry comprising two copies of the DBD and one DNA duplex, although the latter was truncated from a 22mer to a 16mer. This yielded a good solution and, in common with the template structure, the DNA formed a pseudo-continuous filament spanning the crystal due to base-pair stacking between DNA fragments in adjacent ASUs. However, there was only sufficient space to accommodate 15 bp per ASU within this filament. For the time being, the DNA model was truncated to the central 14 bp *NBS* site in COOT^26^ before real space refining using “chain refine”. The model was subsequently refined with REFMAC5^27^, using jelly body refinement giving *R*_work_ and *R*_free_ values of 0.363 and 0.404, respectively, to 2.9 Å resolution. Inspection of the electron density at this stage revealed evidence for the missing N-terminal domains (NTDs). A template for these was generated using SCULPTOR^28^ from the *Geobacillus thermoleovorans* NOC structure (PDB accession code 7NFU)^6^, where the corresponding domain shares 67% sequence identity with *B. subtilis*. After quickly tidying the output of the REFMAC5 job in COOT, this was put back into PHASER as a search model together with two copies of the NTD template. However, PHASER was only able to place one of the latter sensibly. After further jelly body refinement of this partial model (giving *R*_work_ and *R*_free_ values of 0.313 and 0.351, respectively, to 2.9 Å resolution) the electron density was inspected again in COOT, at which point it was possible to manually dock the missing domain into fragmented density. Following restrained refinement in REFMAC5, the density for the DNA was much clearer, enabling the missing DNA bases to be fitted. In one strand the 5’ base was flipped out, and in the other, the 3’ base was flipped out, enabling a sticky-ended interaction (with a one-base overhang) between the duplexes in adjacent ASUs. After further iterations of model building in COOT and restrained refinement in REFMAC5, the final model was produced with *R*_work_ and *R*_free_ values of 0.230 and 0.277, respectively, to 2.9 Å resolution. Refinement and validation statistics are summarized in Table 1.

---

**Supplementary Figure 1.**
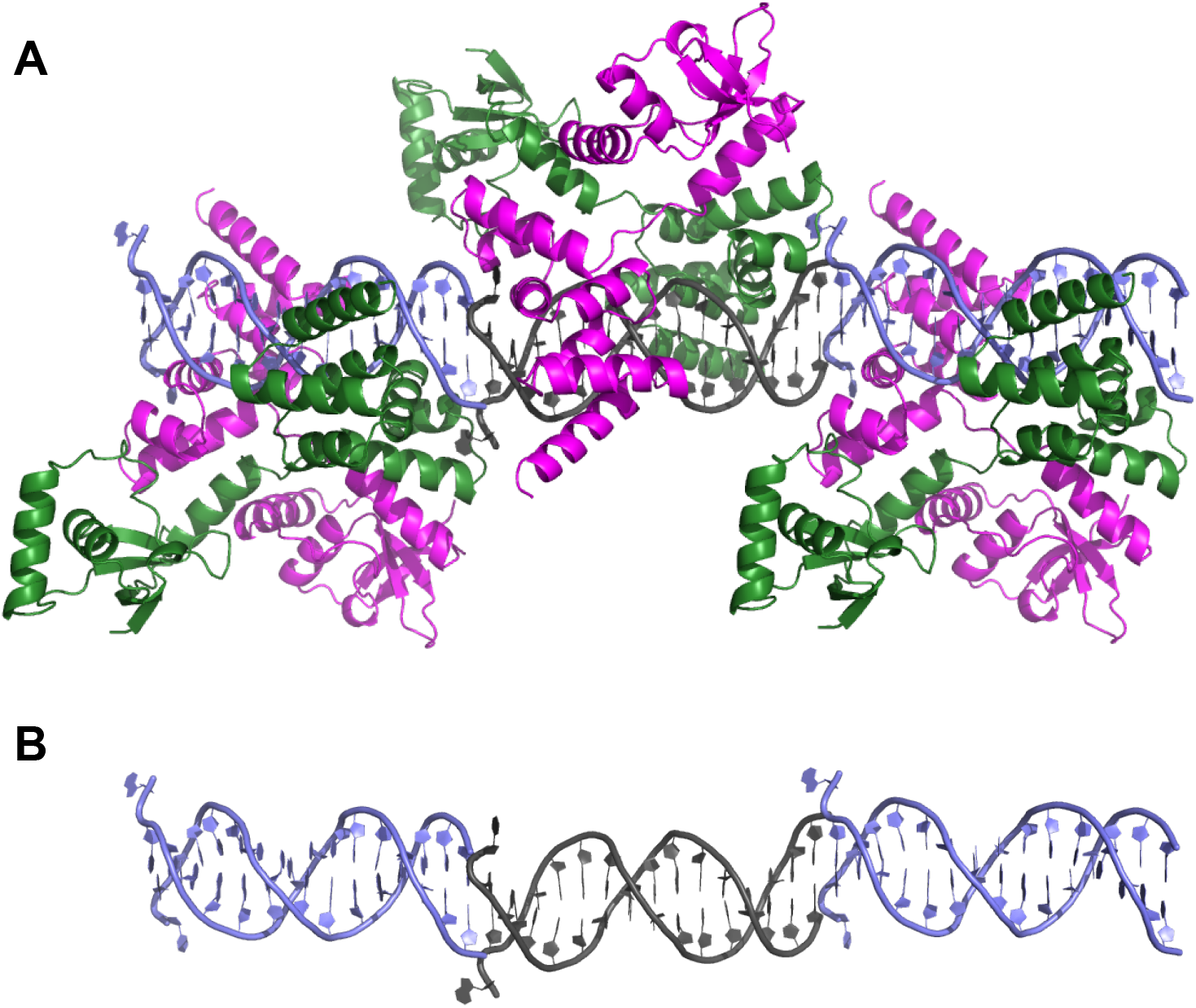
**(A)** Co-crystal structure of *B. subtilis* NocΔCTD-*NBS* complex in the context of the crystal. Neighboring DNA duplexes (light blue and grey) stack and interact to form pseudo-continuous DNA filaments running through the crystal. **(B)** Only the *NBS* DNA portion from the NocΔCTD-*NBS* complex is shown for clarity. Note the flipped-out bases at the junctions between the duplexes.

**Supplementary Figure 2.**
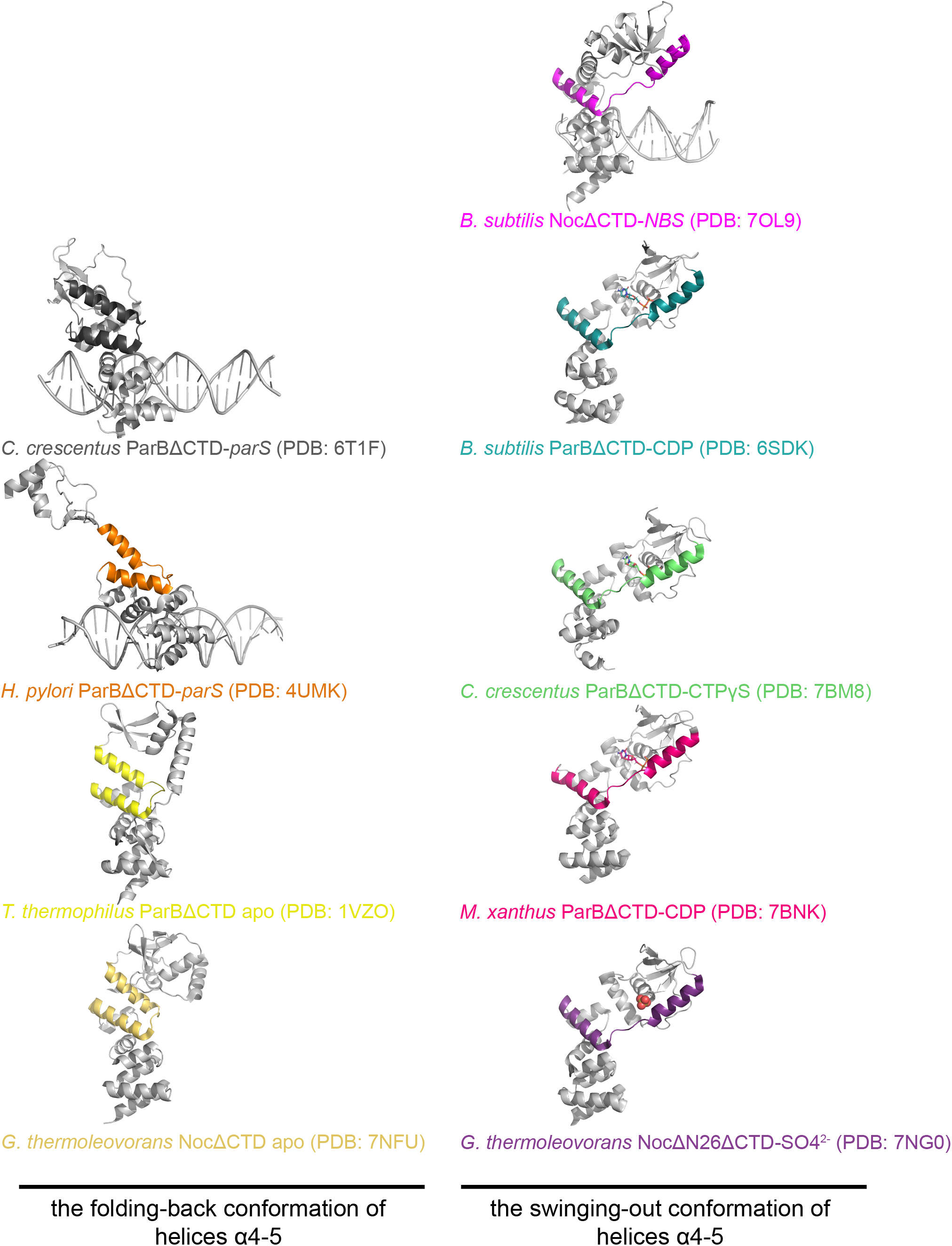
Structures of available chromosomal ParB and Noc, either in the apo-, DNA-bound, or nucleotide-bound states. Only the pair of helices α4 and α5 are highlighted in bright colors to show either the folding-back (left column) or the swinging-out conformation (right column).

**Supplementary Figure 3.**
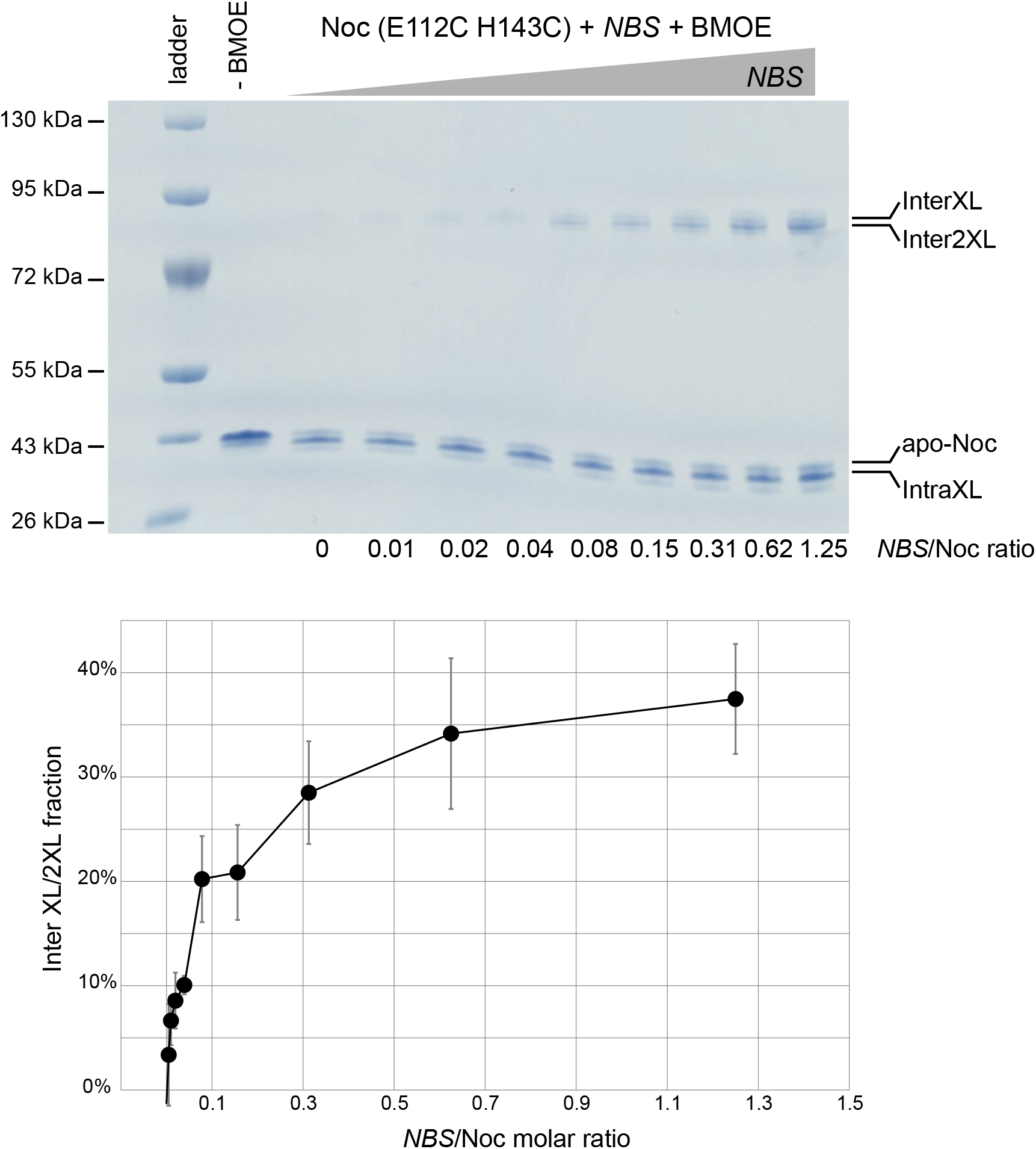
Crosslinking of *B. subtilis* Noc (E112C H143C) in the presence of an increasing concentration of *NBS* and absence of CTP. Crosslinked species were resolved on an acrylamide gel and stained with Coomassie, and InterXL/2XL fractions were quantified. Each crosslinking experiment was run in triplicate.

